# The Role of Left Dorsolateral Prefrontal Cortex in Language Processing

**DOI:** 10.1101/230557

**Authors:** Jana Klaus, Dennis J. L. G. Schutter

## Abstract

In addition to the role of left frontotemporal areas in language processing, there is increasing evidence that language comprehension and production require control and working memory resources involving the left dorsolateral prefrontal cortex (DLPFC). The aim of this study was to investigate the role of the left DLPFC in both language comprehension and production. In a double-blind, sham-controlled crossover experiment, thirty-two participants received cathodal or sham transcranial direct current stimulation (tDCS) to the left DLPFC while performing a language comprehension and a language production task. Results showed that cathodal tDCS increases reaction times in the language comprehension task, but decreases naming latencies in the language production task. However, additional analyses revealed that the polarity of tDCS effects was highly correlated across tasks, implying differential individual susceptibility to the effect of tDCS within participants. Overall, our findings demonstrate that left DLPFC is part of the complex cortical network associated with language processing.

---

Language comprehension and speech production are unique human abilities. To what extent these abilities recruit shared cortical regions of the left frontotemporal language network has been the primary focus of neuroimaging studies (e.g., Humphreys & Gennari, 2014; Menenti, Gierhan, Segaert, & Hagoort, 2011; Segaert, Menenti, Weber, Petersson, & Hagoort, 2012; Silbert, Honey, Simony, Poeppel, & Hasson, 2014). As a consequence, contributions of regions outside this established cortical network involved in language comprehension and production have largely been neglected (Indefrey and Levelt, 2004; Hickok and Poeppel, 2007; Indefrey, 2011; Price et al., 2011; Pickering and Garrod, 2013). Despite the frontotemporal cortico-centred theories of language processing, there is increasing evidence that other regions contribute as well. According to the Memory-Unification-Control (MUC) model (Hagoort, 2013, 2016), a control mechanism in language processing is located in the left dorsolateral prefrontal cortex (DLPFC). The MUC model is supported by functional magnetic resonance imaging (fMRI) studies reporting activation of the left DLPFC during sentence comprehension (Hashimoto and Sakai, 2002; Cooke et al., 2006; Makuuchi et al., 2009; Stephens et al., 2010; Hsu et al., 2017) and sentence production (Humphreys and Gennari, 2014).

To examine the functional nature of the neural activation patterns more directly, transcranial direct current stimulation (tDCS) is used to directly influence the cortical areas during language processing (Joyal & Fecteau, 2016; Price, McAdams, Grossman, & Hamilton, 2015; cf. Westwood & Romani, 2017). Yet, the number of studies that have targeted the left DLPFC is limited. For language comprehension tasks, both anodal and cathodal tDCS over the left DLPFC have been found to improve performance in the comprehension of idioms (Sela et al., 2012; Mitchell et al., 2016) and garden-path sentences (Hussey et al., 2015). For language production, anodal tDCS over the left DLPFC has been reported to decrease naming latencies in action and object naming (Fertonani et al., 2010, 2014), error rates in a scene description task (Nozari et al., 2014a), and a semantic interference effect during picture naming (Wirth et al., 2011). However, it is difficult to infer from previous studies whether both language comprehension and production recruit the same control system, primarily because these two faculties so far have only been tested between participants and studies, and because both the employed tasks and the experimental parameters differed substantially. Thus, while there is increasing evidence from single studies that the left DLPFC is involved in both language comprehension and production, a direct comparison of these two processes is still lacking.

Here we investigated the effects of cathodal tDCS on language processing performance. Specifically, we examined the involvement of the left DLPFC in a picture-mediated language comprehension and production task. Additionally, the effect of cathodal tDCS on task difficulty was examined by manipulating task demands. If these abilities recruit the DLPFC as a critical control region outside of the frontotemporal language network, we predicted worse performance (i.e., increased reaction times and/or higher error rates) during cathodal tDCS compared to sham as an indicator of the involvement of the DLPFC in language processing. If, by contrast, language processing proceeds independently of working memory and control processes associated with the DLPFC, tDCS will not cause any behavioral changes.

## Methods

### Participants

Thirty-two healthy volunteers (22 female; mean age: 22.9 years, *SD* = 2.6, range: 19 – 28) participated in the study. All were native Dutch speakers, right-handed (measured by the Edinburgh Inventory of Handedness, *M* = 93.9 %, *SD* = 5.2, range: 83.3-100.0; Oldfield, 1971), and had normal or corrected-to-normal vision. None of them reported a history of neurological or psychiatric illnesses, current pregnancy, drug or alcohol addiction, skin diseases or allergies, metallic objects in their heads or any type of stimulator in their body, or family history of epilepsy. Participants gave written informed consent prior to the study, which was approved by the medical ethics committee of the Radboud University Medical Centre in Nijmegen. They received 25 € in exchange for their participation.

### Tasks

Figure 1 illustrates two example trials of the three different tasks used in the current experiment.

**Figure 1.**
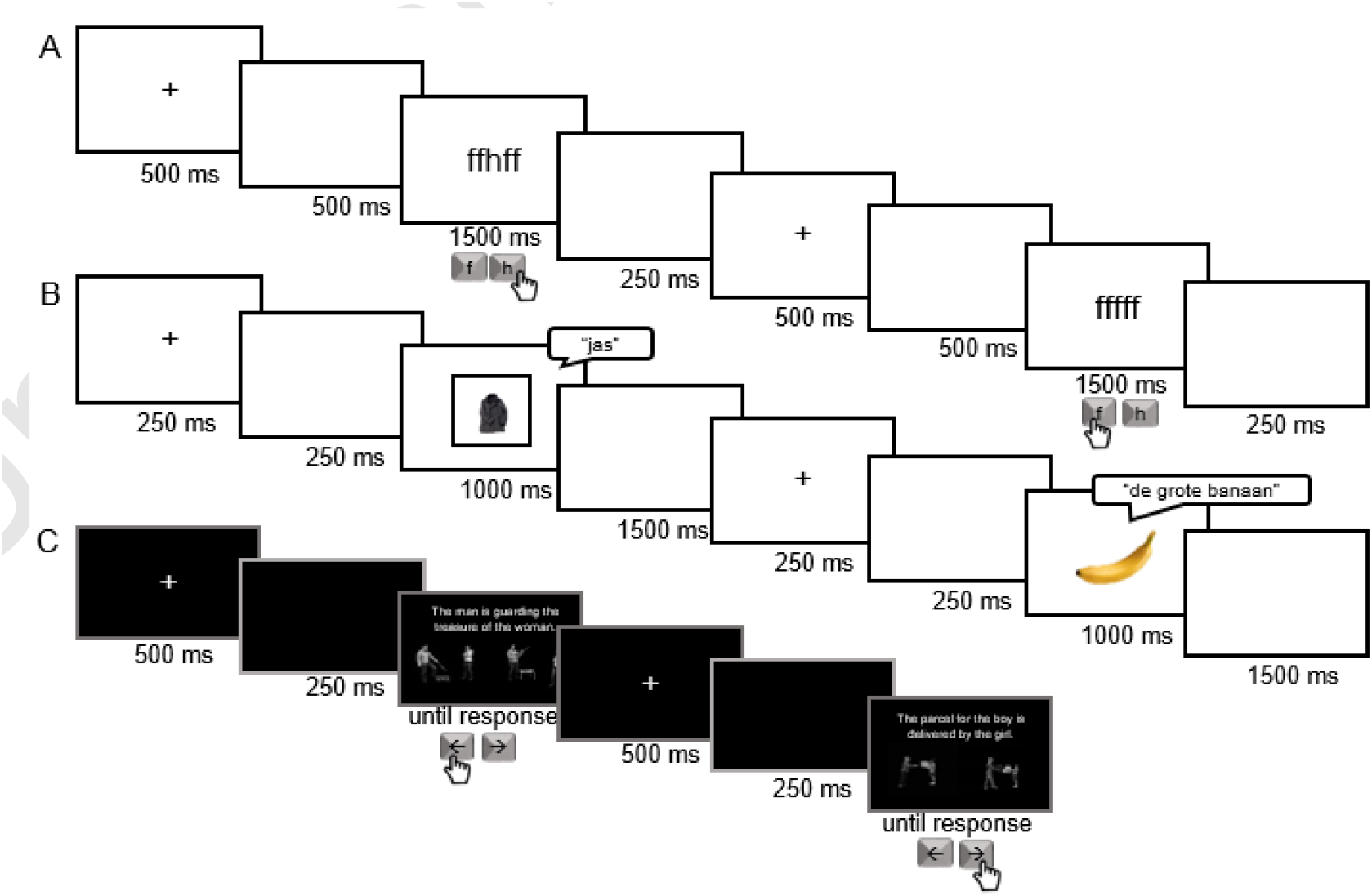
Illustration of two example trials each and the respective required response for the flanker task (A), the language production task (B), and language comprehension task (C).

#### Language production task

112 coloured photographs of everyday objects were chosen as stimuli. All objects could be named with a Dutch mono-or bisyllabic noun and were matched for log frequency (*M* = 2.63, *SD* = 0.52, range: 1.18-3.56) in the SUBTLEX-NL database (Keuleers et al., 2010). The pictures eliciting a bare noun utterance were scaled to fit a black frame of 360 x 360 pixels. For the complex noun phrase condition, the size of the pictures was tripled (1063 x 1063 pixels). Participants were instructed to name the pictures as quickly and correctly as possible, using a bare noun (e.g., “tafel” [table]) or a complex noun phrase (e.g., “de grote tafel” [the big table]) depending on whether the presented stimulus fit into the accompanying frame or not. Note that in the complex noun phrase condition, speakers had to choose between two different grammatical genders before initiating the response (“de” [masculine/feminine] or “het” [neuter] in Dutch), while the following adjective was identical for all items (“grote” [big]).

Utterance format was tested within items and participants but between sessions, e.g. in one session, participants named a given picture as a bare noun and in the other session as a complex noun phrase. Eight pseudorandomised lists were created for the experimental blocks, accounting for the constraints that no more than three consecutive trials required the same utterance format, items from the same semantic category were separated by at least five intervening trials, and items with the same phonological onset were separated by at least four intervening trials. Participants received different lists in the two sessions. Each experimental block consisted of 112 trials. The task was administered in Presentation software (Version 18.1, Neurobehavioral Systems, Inc., Berkeley, CA, www.neurobs.com). Naming latencies were measured to the closest millisecond through a voice-key connected to the microphone placed in front of the participant. Naming errors were coded online by the experimenter.

#### Language comprehension task

The comprehension task consisted of matching a visually presented sentence to one of two picture stimuli. The pictures were depictions of eight different transitive actions which illustrated a subject (i.e., agent) performing an action on a direct object for an indirect object (i.e., patient) (e.g., a boy delivering a parcel to a girl; Menenti, Gierhan, Segaert, & Hagoort, 2011; Segaert, Menenti, Weber, & Hagoort, 2011). Importantly, the two pictures were always completely identical, except that the direct object was exchanged, but belonged to the same associative-semantic category to increase lexical competition (e.g., when the target sentence was “de jongen bezorgt het pakket voor het meisje” [The boy is delivering the parcel to the girl] the pictures of a boy delivering a parcel and of a boy delivering a letter were presented). Additionally, the difficulty of the target sentence was varied such that it either appeared in active or passive voice. The target sentence was presented in the upper half of the screen and the two pictures next to each other in the lower half of the screen. The position of the target picture (left or right) was counterbalanced within blocks.

Eight pseudorandomised lists were created for the experimental blocks, accounting for the constraints that no more than three consecutive trials were presented in the same voice (active vs. passive), pictures of the same action were separated by at least three intervening trials, and a target picture appeared for no more than three consecutive trials in the same position of the screen (left vs. right). Participants received different lists in the two sessions. Each experimental block consisted of 112 trials. The task was administered in Presentation software (Version 18.1, Neurobehavioral Systems, Inc., Berkeley, CA, www.neurobs.com), and responses were given through the keyboard placed in front of the participant.

#### Flanker task

This task was implemented as a control task to verify the sensitivity of the current stimulation protocol. Nozari, Woodard, & Thompson-Schill (2014) reported increased reaction times in a flanker task during cathodal tDCS over the left DLPFC, indicating a crucial involvement of this region in cognitive control and response inhibition. Replicating these results would provide evidence that in the current study, we also successfully targeted this region.

A modified version of the browser-based flanker task provided by the Experiment Factory (Sochat et al., 2016) was used as the experimental task in both sessions. A string of five letters consisting of f’s and h’s was presented in the centre of the screen, and participants were asked to respond to the middle letter by pressing the appropriate key (f or h) on the keyboard. Half of the stimuli were congruent (i.e. the middle letter was identical to the flanking letters, ‘fffff’ or ‘hhhhh’) and the other half incongruent (i.e. the middle letter differed from the flanking letters, ‘ffhff’ or ‘hhfhh’). Stimulus conditions were generated randomly for each participant. Each experimental block consisted of 100 trials. The task was administered in Google Chrome, and responses were made through the keyboard located in front of the participant.

### Transcranial direct current stimulation

Stimulation was delivered in a randomized double-blind fashion by a battery-driven stimulator via two electrodes sponges covered in conductive gel (3×5 cm each; NeuroConn GmbH, Ilmenau, Germany) which were placed under an EEG cap. Cathodal tDCS was delivered by a cathodal electrode over left LPFC (placed between F3 and F7) and the anodal electrode positioned anterior to the vertex (between Fz and Cz) (Figure 2). Following a 30-second ramp up, tDCS was delivered at 2 mA intensity (current density: 0.133 mA/cm^2^), and continued throughout the experimental tasks (i.e., online). Depending on the length of breaks between tasks required by each participant, stimulation duration varied between participants between 20 to 30 minutes. Impedance of the electrodes was below 15 kΩ during stimulation. Real and sham stimulation were randomly assigned across the two sessions, with half of the participants receiving real tDCS in the first session and sham tDCS in the second session, and the other half sham tDCS in the first session and real tDCS in the second session. Experimenter blinding was achieved by using a pre-assigned code entered into the DC stimulator at the beginning of each session.

**Figure 2.**
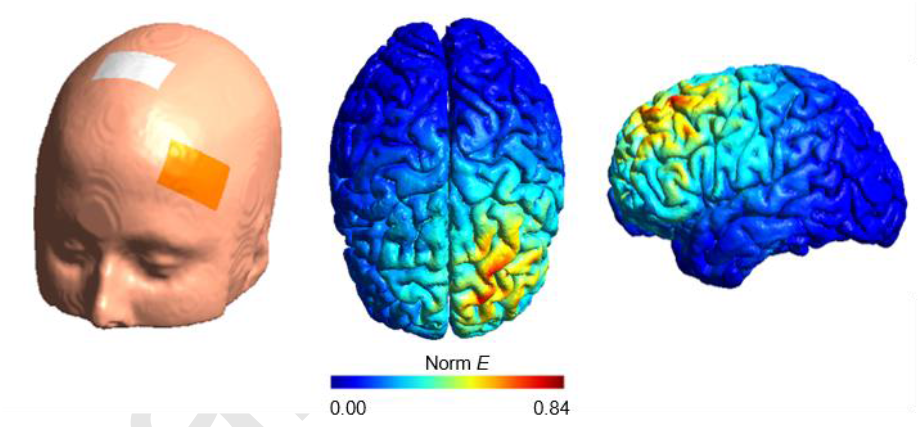
A simulation was performed on a standard brain to estimate the electric field density and distribution using SimNIBS (Opitz et al., 2015). The cathodal electrode was placed over the left DLPFC (between F3 and F7), and the reference electrode anterior to the vertex (between Fz and Cz).

### Procedure

Participants were tested in two sessions of approximately 45 minutes each. Each session was separated by at least 48 hours to minimize carry-over effects, and took place at the same time of the day. Prior to the experiment, participants received written and oral information about the study, after which they were asked to fill in the consent and screening forms. Afterwards, tDCS was administered, during which participants performed the experimental tasks, for which instructions were presented on the screen. Each task was preceded by a short practice block familiarising the participants with the procedure. Participants always started with the flanker task as the control task, and the order of the comprehension and production task was counterbalanced across participants and sessions. After each session, participants received a form in which they were asked to indicate any subjectively experienced side effects. At the end of the second session, participants were debriefed about the purpose of the study.

### Data reduction and analysis

Trials in which a wrong response was given (i.e., a wrong button press or a wrong/disfluent utterance) were discarded from the reaction time analyses (production: 9.7%; comprehension: 3.0%; flanker: 4.1%). Observations deviating from a participant’s median by more than three standard deviations, computed separately for cathodal and sham tDCS, were marked as outliers and also removed (production: 1.6%; comprehension: 1.7%; flanker: 1.5%). Additionally, in the production task the item “garde” (whisk) was removed from further analyses because of a mean error rate larger than 30%.

Statistical analyses were computed with generalised mixed-effects models (GLMEMs) using the *lme4* package (version 1.1.12, Bates, Mächler, Bolker, & Walker, 2015) in R (version 3.3.3, R Core Team, 2017). Contrary to linear mixed effects models, GLMEMs can account for the right-skewed shape of the RT distribution without the need to transform and standardise the raw data (Lo and Andrews, 2015). For the reaction time data, we fitted an identity function to reaction time data assuming a Gamma distribution (i.e., right-skewed with a long tail in the slow RTs). Error rates were analysed using mixed logit regression (Jaeger, 2008). For all tasks, we included by-participant intercepts to account for interindividual variability in overall task performance, as well as by-participant slopes for the main effect of tDCS condition. Additionally, we included a by-participant and by-item slope for difficulty in the language tasks (i.e., utterance format in the production task and voice in the comprehension task). The alpha level was set to < 0.05 (two-tailed) for all analyses. Tasks and tDCS condition were fully crossed and tested within participants to allow for a direct comparison of the involvement of DLPFC in individual language processing.

## Results

Participants tolerated tDCS well and only reported a slight tingling or itching under the electrodes during the ramp-up phase. A Chi-square test comparing the coded post-session responses asking about perceived side effects revealed no difference between real and sham stimulation (*χ*^2^(1) = 0.58, *p* = .445), indicating that blinding was successful.

Figures 3 and 4 display the reaction times and error rates, respectively, for all three tasks, broken down by tDCS condition (cathodal vs. sham) and the respective within-task difficulty factor (flanker: stimulus congruency [congruent vs. incongruent]; production: utterance format [bare noun vs. complex noun phrase]; comprehension: voice [active vs. passive]).

**Figure 3.**
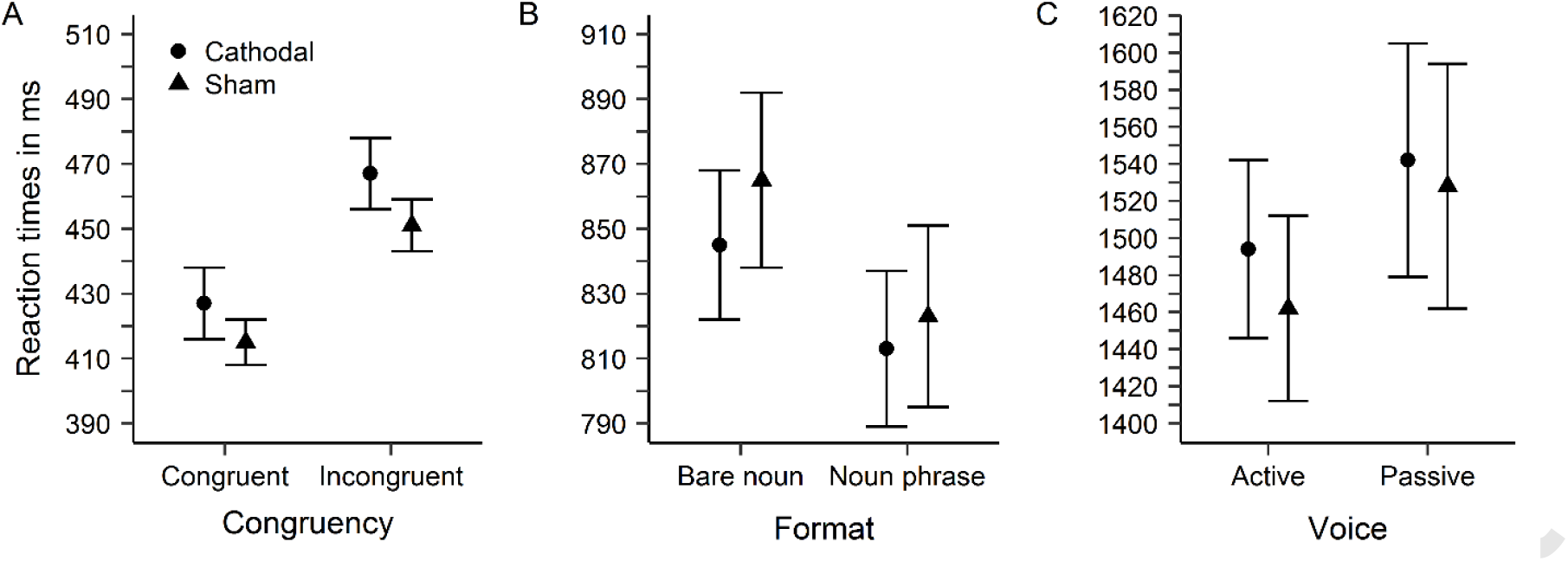
Mean reaction times (± SEM) for the three different tasks, aggregated across participants and broken down by tDCS type (cathodal vs. sham) and task difficulty. A: Flanker task by stimulus congruency (congruent vs. incongruent). B: Language production task by utterance format (bare noun vs. complex noun phrase). C: Language comprehension task by voice (active vs. passive).

**Figure 4.**
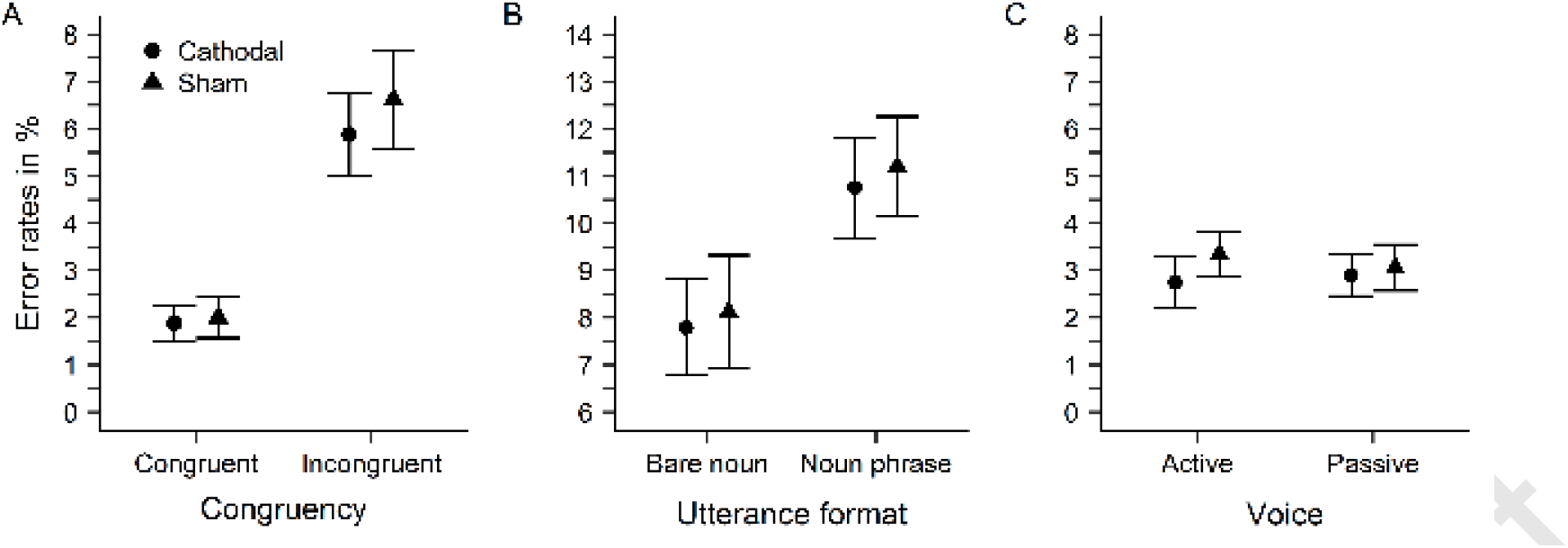
Mean error rates (± SEM) for the three different tasks, aggregated across participants and broken down by tDCS type (cathodal vs. sham) and task difficulty. A: Flanker task by stimulus congruency (congruent vs. incongruent). B: Language production task by utterance format (bare noun vs. complex noun phrase). C: Language comprehension task by voice (active vs. passive).

### Flanker task

Participants responded faster and with fewer errors to congruent compared to incongruent stimuli, demonstrating the classic flanker effect (RTs: β = -18.58, *SE* = 0.77, *t* = -24.27, *p* < .0001; errors: β = -0.62, *SE* = 0.07, *z* = -8.48, *p* < .0001). Cathodal tDCS increased reaction times compared to sham tDCS (β = 9.14, *SE* = 3.41, *t* = 2.68, *p* = .007), but did not affect error rates (β = -0.06, *SE* = 0.10, *z* = -0.58, *p* = .821). The size of the congruency effect did not differ between tDCS conditions (*ps* > .372). These results show that the current stimulation montage was successful in decreasing performance in a task recruiting the left DLPFC (see Nozari, Woodard, et al., 2014).

### Language production

Naming latencies were shorter during cathodal compared to sham tDCS (β = -11.80, *SE* = 4.59, *t* = -2.57, *p* = .010), while error rates were not affected (β = -0.02, *SE* = 0.06, z = -0.35, *p* = .726). Participants were faster, but made more errors when producing complex noun phrases compared to bare nouns (naming latencies: β = 20.30, *SE* = 5.10, *t* = 3.98, *p* < .0001; error rates: β = -0.23, *SE* = 0.07, *z* = -3.55, *p* < .0001). No effect of tDCS was observed on utterance format (*ps* > .233).

### Language comprehension

Reaction times were slower during cathodal compared to sham tDCS (β = 9.34, *SE* = 4.67, *t* = 2.00, *p* = .045) and faster in response to active sentences compared to passive sentences (β = -23.88, *SE* = 5.67, *t* = -4.20, *p* < .0001). The interaction of tDCS and difficulty was not significant (*p* > .512), and also no significant effects were found in the error rate analysis (*ps* > .380).

### Individual effects of tDCS

The magnitude of the behavioral effects of tDCS is subject to substantial inter-individual variability (Wiethoff et al., 2014; Chew et al., 2015; Li et al., 2015). The within-participant design of the current study allowed us to directly assess these possible variations. To obtain individual measures of tDCS efficacy, we correlated the individual tDCS effects for each task (i.e., RT_cathodal_ – RT_sham_). As can be seen in Figure 5, the individual effects were highly, and exclusively positively, correlated across tasks, suggesting that the behavioural outcome induced by tDCS is consistent within participants.

**Figure 5.**
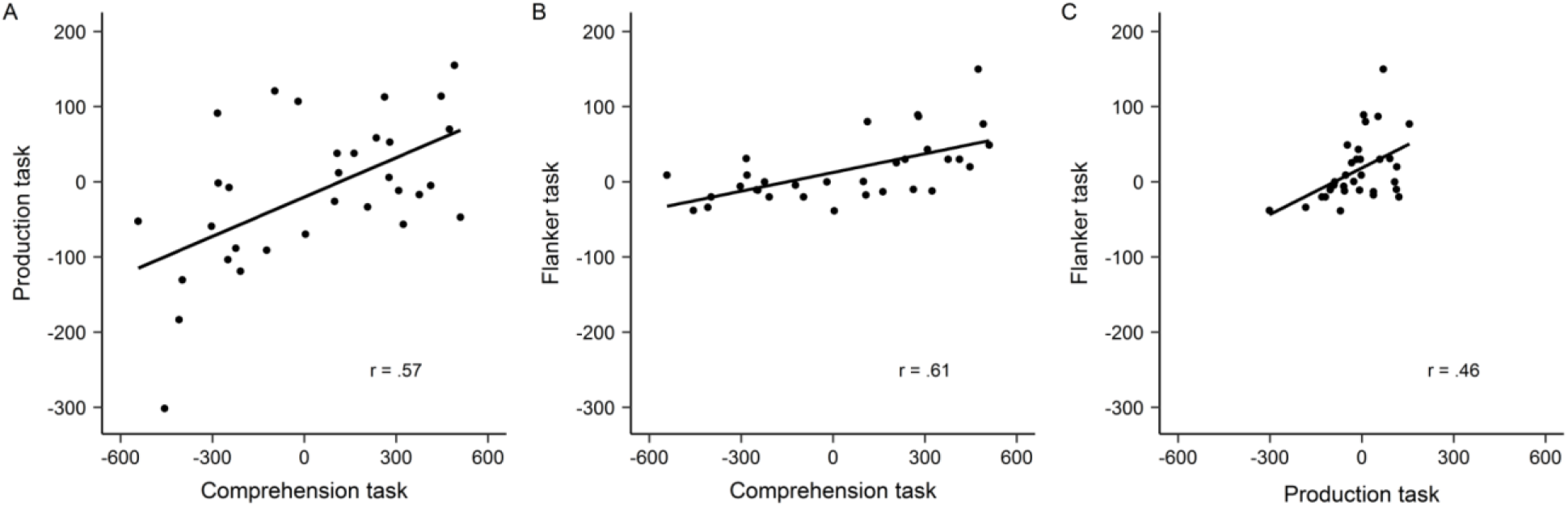
Correlation plots of individual tDCS effects (RT_cathodal_ – RT_sham_), displaying positive correlations across tasks.

### Effects across sessions

Finally, it should be noted that the within-participant design of the current study also comes at a cost given the low test-retest reliability reported in previous studies (Horvath et al., 2016; Wörsching et al., 2017). The application of tDCS (real vs. sham) was counterbalanced across participants. Consequently, half of the participants received cathodal tDCS in the second session, at which point they were already familiar with the tasks and subsequently, practice effects may have obstructed the overall effects of tDCS. Indeed, when including the factor session (first vs. second) in the analyses reported above, we found significant interactions of tDCS condition and session for all tasks (production: β = -30.06, *SE* = 5.69, *t* = -5.28, *p* < .0001; comprehension: β = -32.64, *SE* = 7.18, *t* = -4.50, *p* < .0001; flanker: β = 13.70, *SE* = 5.07, *t* = 2.70, *p* = .007). Therefore, we reanalysed the data for the first session only, in which all participants were equally naïve to the tasks, but tDCS was tested between participants. For all tasks, we still found a main effect of tDCS (production: β = 42.21, *SE* = 17.28, *t* = -2.44, *p* = .015; comprehension: β = -16.65, *SE* = 7.92, *t* = -2.28, *p* = .022; flanker: β = 22.00, *SE* = 6.00, *t* = 3.67, *p* < .001), with higher reaction times for the cathodal group in the comprehension and flanker task, but lower naming latencies in the production task compared to sham. This analysis not only confirms the reliability of our overall results, but also shows that task repetition may induce practice effects and influence the effects of tDCS in the second session.

## Discussion

The present study showed that, on a global level, cathodal tDCS increased reaction times in the comprehension task, but decreased naming latencies in the production task compared to sham tDCS. Thus, our results suggest involvement of the left DLPFC in both language comprehension and production, although suppressing activity in this region appears to have differential effects on production and comprehension, respectively.

The overall disruptive effect of cathodal tDCS on language comprehension performance may be explained in terms of an involvement of working memory, which has been related to DLPFC activity (Mottaghy et al., 2000; Funahashi, 2006; Brunoni and Vanderhasselt, 2014; D’Esposito and Postle, 2015; Mansouri et al., 2015), in this process. This is in line with behavioural evidence linking these two processes (Daneman and Carpenter, 1980; King and Just, 1991; Just and Carpenter, 1992; Caplan and Waters, 1999), and previous tDCS studies which reported an involvement of left DLPFC in reading garden-path sentences (facilitation from anodal tDCS; Hussey, Ward, Christianson, & Kramer, 2015) and idioms (facilitation from anodal and cathodal tDCS; Sela et al., 2012; Mitchell et al., 2016). In the current study, participants had to keep the contents of the presented sentence activated while matching them to one of two pictures, and performance in this task was worse if the left DLPFC was disrupted. This implies that mapping the syntactic features of a sentence onto visual input to successfully comprehend the sentence may critically rely on working memory. Notably, the current study is the first to report an inhibitory effect of cathodal tDCS across the left DLPFC in comprehension. We show that matching both active and passive voice sentences to one of two pictures is significantly impaired during the administration of cathodal tDCS to the left DLPFC.

Another possibility is that the performance in language comprehension was influenced by changes in motivational control. According to frontal lateralization theories (e.g., Schutter & Harmon-Jones, 2013; Spielberg et al., 2011), the left DLPFC is linked to approach-motivation, and increased activity in this region may influence cognitive functions by way of modifying mental effort (Harmon-Jones et al., 2012; Schutter, 2014). Disrupting cortical excitability of the left DLPFC by cathodal tDCS may have resulted in a decrease in approach-related motivation and mental effort necessary for executing a complicated task like sentence comprehension.

In the language production task, cathodal tDCS decreased naming latencies compared to sham tDCS. Crucially, the effect of tDCS (i.e., shorter naming latencies during cathodal tDCS compared to sham) did not differ between utterance formats. This suggests that for both utterance formats, suppressing cortical excitability of the left DLPFC aided language production. However, this finding is at odds with our prediction, as we would have expected a disruptive effect of cathodal tDCS in the case of DLPFC involvement in production (i.e., analogous to the language comprehension task). Yet, it is not unreasonable to assume that interfering with the activity of the left DLPFC during a highly automated task such as picture naming might actually facilitate lexical retrieval. That is, because naming required either only the retrieval of a single word or of a highly predictable utterance in the current task, control demands admittedly were not very high. Decreasing activity of the DLPFC as an instance controlling utterance preparation may have fine-tuned the picture naming process such that fewer computations were needed for successful production. A critical test then would be to increase the overall difficulty of the naming task, for instance by extending the paradigm to a sentence production task. We assume that in such a case, a higher load on working memory and cognitive control will reveal the disruptive effect of cathodal tDCS.

An important finding from our study is that the global effects described above varied substantially between, but were consistent within individuals. That is, when looking at the effects on the individual level, we could show that the direction of the effects induced by tDCS (i.e., facilitatory or inhibitory) was highly correlated across tasks. Thus, despite the large distribution of effects both with respect to their magnitude and polarity, their directionality (i.e., whether tDCS inhibited or facilitated task performance) was consistent across tasks. This is especially relevant with regard to the counterintuitive overall facilitation effect we found in the language production task: Nine out of thirty-two participants showed an RT increase larger than 50 ms, suggesting that production was affected differently by inhibiting the left DLPFC. While this interpretation is highly speculative and requires further experimental testing, it is possible that these participants generally rely more on working memory resources during language processing, which caused a larger performance decrement in the cathodal session. By contrast, participants exhibiting a large facilitation effect in the language production task also tended to benefit from cathodal tDCS in the language comprehension task. This finding can be interpreted in two ways: One possibility is a differential involvement of the left DLPFC in language processing between participants, with some participants relying on this cortical region more than others. Alternatively, it is possible that the physiological response to tDCS differs between individuals, causing variability in the behavioural response. This is a promising outlook for future research, as the current data suggest that individual differences in DLPFC recruitment and/or response to tDCS differentially affect the behavioural outcome in language processing.

Three issues need to be addressed. First, it should be noted that the variability in reaction times was large in the comprehension task because unlike for the other tasks, stimuli remained on screen until participants made a response, but there was no response deadline. We selected this procedure because Manenti et al. (2008) had found higher reaction times following repetitive transcranial magnetic stimulation (rTMS) to the left DLPFC in a similar experimental setup. Our study replicates these findings using cathodal tDCS, suggesting that inhibitory effects of tDCS do not depend on fixed time intervals, but perhaps also arise when participants self-pace their responses.

Second, the finding that complex noun phrases were produced faster than bare nouns was unexpected. Typically, the production of complex noun phrases results in longer naming latencies, as they necessitate planning more elements prior to articulation (e.g., Bürki, Sadat, Dubarry, & Alario, 2016; Jescheniak, Schriefers, & Hantsch, 2003). However, since noun phrase production in the current study always required the production of the determiner (“de” or “het”) and the same adjective (“grote” [big]), its structure was highly predictable, encouraging the strategy to quickly utter these two elements while planning the rest of the utterance online. Note that during noun phrase production, error rates, reflected mostly in disfluencies and utterance repairs, were substantially higher compared to bare noun production, suggesting a speed-accuracy trade-off in these conditions. Thus, when taking into account both naming latencies and error rates, we would argue that the complex noun phrase production still was the more difficult condition, while bare noun production reflected more “pure” picture naming. More importantly for the current study, however, tDCS did not affect naming latencies differentially as a function of utterance format. Thus, regardless of what caused the faster naming latencies in the complex noun phrase condition, effects of this manipulation were not modulated by functioning of the left DLPFC.

Third, previous research has shown that applying 2 mA cathodal tDCS across the motor cortex can result in excitatory rather than inhibitory activity after stimulating for 20 minutes (Batsikadze et al., 2013). In the current study, tDCS was applied online (i.e., during execution of the task). Given that the tasks combined took between 20 and 30 minutes, we cannot rule out the possibility that initial inhibitory effects interacted with later excitatory effects. We did, however, take available precautions to control for this possible confound (i.e., task order was counterbalanced across participants and stimulation sessions), so we do not think this finding from the motor cortex had a systematic effect on our results. Nevertheless, we acknowledge that it may have added some noise to the data, and certainly raises interesting research questions for further studies.

In conclusion, our findings support the MUC model by showing evidence for involvement of the left DLPFC in language production and comprehension. Additional research is needed to further examine the origins of the interindividual differences in the polarity-dependent effects of tDCS on behaviour, and the specific role of the left DLPFC in language processing.

## Acknowledgments

This work was supported by the German Research Council under Grant KL 2933/2. We thank Katrien Segaert for providing the stimulus materials used in the language comprehension task.

